# Vimo: Visual Analysis of Neuronal Connectivity Motifs

**DOI:** 10.1101/2022.12.09.519772

**Authors:** Jakob Troidl, Simon Warchol, Jinhan Choi, Jordan Matelsky, Nagaraju Dhanyasi, Xueying Wang, Brock Wester, Donglai Wei, Jeff W. Lichtman, Hanspeter Pfister, Johanna Beyer

**Affiliations:** School of Engineering & Applied Sciences, Harvard University; Department of Cellular & Molecular Biology, Harvard University; Applied Physics Laboratory, Johns Hopkins University; Department of Bioengineering, University of Pennsylvania; Department of Computer Science, Boston College

**Keywords:** Visual motif analysis, Focus&Context, Scientific visualization, Neuroscience, Connectomics

## Abstract

Recent advances in high-resolution connectomics provide researchers with access to accurate petascale reconstructions of neuronal circuits and brain networks for the first time. Neuroscientists are analyzing these networks to better understand information processing in the brain. In particular, scientists are interested in identifying specific small network motifs, i.e., repeating subgraphs of the larger brain network that are believed to be neuronal building blocks. Although such motifs are typically small (e.g., 2 - 6 neurons), the vast data sizes and intricate data complexity present significant challenges to the search and analysis process. To analyze these motifs, it is crucial to review instances of a motif in the brain network and then map the graph structure to detailed 3D reconstructions of the involved neurons and synapses. We present *Vimo*, an interactive visual approach to analyze neuronal motifs and motif chains in large brain networks. Experts can sketch network motifs intuitively in a visual interface and specify structural properties of the involved neurons and synapses to query large connectomics datasets. Motif instances (MIs) can be explored in high-resolution 3D renderings. To simplify the analysis of MIs, we designed a continuous focus&context metaphor inspired by visual abstractions. This allows users to transition from a highly-detailed rendering of the anatomical structure to views that emphasize the underlying motif structure and synaptic connectivity. Furthermore, *Vimo* supports the identification of motif chains where a motif is used repeatedly (e.g., 2 - 4 times) to form a larger network structure. We evaluate *Vimo* in a user study and an in-depth case study with seven domain experts on motifs in a large connectome of the fruit fly, including more than 21,000 neurons and 20 million synapses. We find that *Vimo* enables hypothesis generation and confirmation through fast analysis iterations and connectivity highlighting.

## 1 Introduction

Recent developments in high-throughput electron microscopy (EM) have allowed large-scale brain mapping at the level of individual synapses, i.e., connectomics. These new brain tissue datasets, along with advances in automated segmentation, enable scientists to accurately reconstruct 3D wiring diagrams of the biological neural networks. For example, the new *H01* dataset [50] captures a cubic millimeter of human brain tissue containing about 57, 000 segmented neurons, and their 150 million synapses, automatically reconstructed from 1.4 petabytes of imaging data. Once the data is fully proofread, studying neuronal connectivity is the key to understanding how the brain computes. Since analyzing an entire brain network is infeasible, neuroscientists use motif analysis as a divide-and-conquer approach to extract small, and comprehensible subgraphs from the brain network, which represent neuronal building blocks [21, 24, 28, 47, 62]. However, synapse-level motif analysis of connectomes is still underexplored. The reason for that is three-fold: First, few datasets exist that are large enough to contain thousands of complete and proofread neurons at an image resolution that resolves individual synapses [12, 34, 44, 50, 54]. Thus, previous approaches mainly focused on small volumes and truncated networks [1] or used heuristics to estimate synaptic connections in lower-resolution data [51]. Second, large graphs are difficult to analyze, especially for non-graph experts. Finding motifs in a large graph is computationally expensive, and most previous motif analysis tools require programming experience. Third, neurons and synapses have a dual representation in the graph compared to the original imaging data. In a graph, neurons are nodes, and synapses are edges. In brain tissue, however, neurons are long, tree-like structures, while synapses are small surfaces. These conceptually different representations make it difficult to visualize how the motif structure relates to a set of connected neurons.

A graph motif is a recurring small subgraph of a larger graph, such as a feedback loop between neurons. For example, researchers have recently identified network motifs in the brain of a fruit fly of three to six neurons that are responsible for context-dependent action selection [24]. However, neuroscientists typically need to combine motif search with an in-depth analysis of motif instances (MIs), which are specific neurons that form a motif. We find that neuroscientists prefer analyzing the 3D anatomy of MIs over abstract connectivity graphs or 2D abstractions since oversimplified 2D layouts can not depict complex spatial arrangments like entangled neurons. Also, 2D layouts can distort neuronal anatomy leading to inaccurate estimates of information flow. Thus, only 3D views provide accurate spatial context for neuroscientists. However, the structure of neurons is so intricate and complex that even for small motifs, a 3D view of several neurons quickly suffers from occlusion and visual clutter. Ideally, a spatial view allows for identifying parts of a neuron actively involved in a motif. Yet, no visual tools exist to explore the 3D nature of neuronal motifs.

In this paper, we present *Vimo*, a novel visualization and analysis tool to explore neuronal motifs and motif chains in large brain networks. Our work makes three main contributions. First, we identify domain-specific goals for neuronal motif analysis based on interviews with neuroscience domain experts. Thus, we derive a set of analysis tasks that are the basis for the design of our web-based visualization tool *Vimo*. Second, we propose a scalable motif analysis workflow, including the design of a flexible *motif searching approach* based on interactive sketching. Scientists draw nodes and edges in a sketching panel to query for their desired motif without having to program. Our queries leverage biological constraints such as neuron types to reduce computational complexity, scale to large brain networks, and give interactive feedback on the number of motif occurrences in the brain network. Additionally, we integrate a *focus&context scheme* for analyzing motif instances and motif-based synaptic chains to facilitate the user’s understanding of how the neuron’s morphology relates to motif connectivity. We do so by de-emphasizing non-motif-relevant parts of the neurons (context) to focus on the motif-relevant aspects. Importantly, we do not change neuron morphology but gradually shift the focus using neuron pruning, exploded views, and hierarchical synapse clustering (see Fig. 1c-e). Third, we demonstrate the integration of our motif analysis workflow into *Vimo*, an open-source [56] and web-based visualization tool. *Vimo*’s novelty lies in combining state-of-the-art methods from scientific visualization, graph theory, and neuroscience into a useful, interactive tool based on a detailed goal and task analysis with domain scientists. We also integrated *Vimo*’s motif sketching interface into the Neuprint connectome analysis platform [7] and released a domain-agnostic UI component for motif sketching [9] to facilitate motif analysis in disciplines other than connectomics. We evaluate *Vimo* with a qualitative user study and a case study of visual input neurons in the fly brain.

**Fig. 1:**
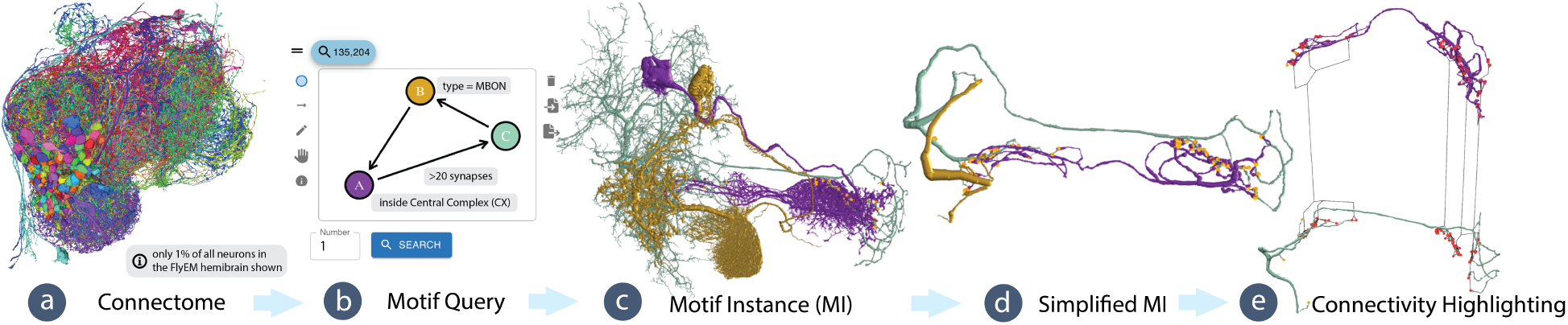
Visual Motif Analysis. (a) In *Vimo*, neuroscientists search large brain networks (view online: [29]) by (b) sketching neuronal connectivity motifs. (c) Neurons forming a motif instance (MI) are visualized in 3D. *Vimo* emphasizes the relationship between the sketched motif and the neurons’ connectivity using a continuous focus&context approach (d, e). First, *Vimo* prunes neuron branches unrelated to the motif (d). Next, users explore the connectivity between neurons in an exploded view that untangles complex neuron morphologies and uses hierarchical synapse clustering and bundling to highlight connections (e). Data: FlyEM Hemibrain [44].

## 2 Related Work

### Connectomics and Motif Analysis

Motif analysis [39] plays a central role in connectome analysis [18, 24, 46, 59]. Hence, numerous computational methods have emerged for examining connectivity motifs within connectomes. Matejek et al. [32] present an optimized and parallelized subgraph enumeration algorithm to count the occurrences of motifs in large brain networks. *DotMotif* [33] is a domain-specific language to write motif queries and compiles to common python tools and the *Cypher* graph query language [17]. *Vimo* builds on *DotMotif*, but offers a visual non-programming interface to create motif queries (see Fig. 2).

**Fig. 2:**
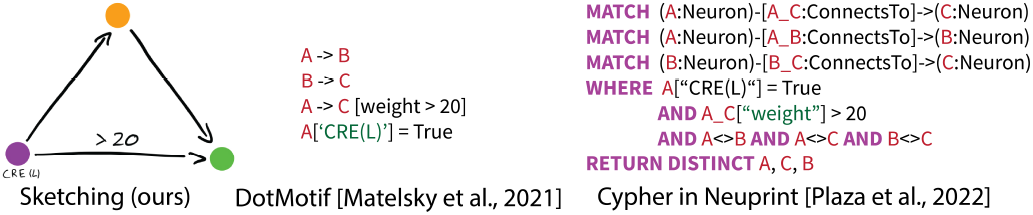
Different motif query approaches. Our visual sketching approach (left), high-level domain-specific language DotMotif [33] (middle), and graph query language Cypher [17] (right).

### Previous Motif Analysis Workflow

In the past, the motif analysis workflow of our collaborators was considerably more laborious due to complex and inaccessible technology. They often cited time constraints and difficult-to-use tools as barriers that prevented them from performing motif analysis. For instance, writing programmatic motif queries is challenging as users must be well-versed with graph query programming languages [17, 40]. Additionally, existing visualization tools have not integrated 3D analysis of MIs into their workflows. Previously, users had to copy neuron IDs obtained from programmatic graph queries into tools like Neuroglancer [30]. Furthermore, none of the available tools offer motif simplification strategies for a more in-depth examination of motif connectivity. Therefore, our collaborators mostly gave up on motif analysis or looked at just very selected motifs in Neuroglancer [30] but did not refine or extend their analyses.

### Visualization for Connectomics

Visual analysis approaches have been used in many subareas of connectomics, including interactive proofreading [13, 20], volume exploration [2], and neighborhood-[55] and morphology analysis [8, 36]. Beyer et al. [3] provide a comprehensive survey of visualization methods in connectomics. Most neural connectivity visualizations use node-link diagrams [2, 41], and therefore do not support a detailed analysis of MIs. Recently, Ganglberger et al. [19] proposed a 2D node-link layout that preserves the spatial context of 3D brain networks. However, their work focuses on macroscale brain parcellations and thus does not extend to nanoscale connectivity. Alternative approaches have used matrix views [22] or visual abstractions in 2D that retain some morphological features, such as neuron branches [1] or arborizations [51]. These methods offer compact visual encodings that allow scientists to get a quick overview of the data. However, the 3D morphology of neurons and their spatial relations in the data is lost. Furthermore, most prior works focus on either very small volumes and, thereby, truncated brain networks [1, 2], hypothetical neuronal circuits [60], or use lower resolution datasets and heuristics to generate synapse positions [51]. Most recently, Plaza et al. [38] developed neuPrint, a tool to query neuronal connectivity quickly in the browser. neuPrint uses the Cypher language [17] to query for motifs but does not focus on visually highlighting the connectivity in MIs.

### Visual Graph Queries

Visual query interfaces have been used in many application areas and allow users to specify queries in an intuitive way [6, 16, 31]. Cuenca et al. [11] propose Vertigo, an approach to construct and suggest graph queries and explore their results in multi-layer networks. Vertigo visualizes all detected subgraphs as an overlayed heatmap to a graph drawing. This approach, however, is not feasible for large connectomic graphs, as all information on the 3D morphology of neurons would be lost. Vigor [37] focuses on effectively summarizing subgraph queries by grouping results based on node features and structural result similarity. In *Vimo*, users can define structural similarity constraints like neuron types directly in the query interface.

### Visualization of Network Motifs

Visualization of motifs has been proposed in different domains. MAVisto [48] supports visual motif analysis in biological networks. However, MIs are only visualized as node-link diagrams, and 3D shapes of biological objects are hidden. Dunne [15] simplifies node-link diagrams of motifs using glyphs. Those simplification techniques do not extend to complex spatial data like neuronal structures. For dynamic networks, Cakmak et al. [5] compare motif significance profiles over time by plotting them as a time series.

### Visual Abstraction of Tree-Like Structures

Different visual abstractions have been proposed to analyze neurons and tree-like structures. Hu et al. [23] project a complex 3D tree-like structure onto a 2D plane while avoiding overlaps. However, their method does not scale to groups of treelike structures, as needed for neuronal motif analysis. Mohammed et al. [36] use a continuous 2D abstraction space to analyze astrocytes and neurons. Neurolines [1] uses a 2D subway map metaphor to display neurons. However, both approaches do not support motif analysis, focus on relatively small subvolumes, and would not visually scale to large connectivity graphs.

## 3 Biological Background

### Brain anatomy

Brain tissue primarily consists of long tubular tree-like *neurons*, which transmit signals to each other via *synapses*. A single neuron can connect to hundreds or thousands of other neurons. The high interconnectivity of neurons combined with their complex morphology results in tangled 3D brain networks that are difficult to analyze (see Fig. 4a). Different neuron types have unique morphological properties and connectivity preferences. Most brains of model organisms, such as the fruit fly (*Drosophila Melanogaster*), are further divided into spatial regions with distinct anatomical and functional properties.

**Fig. 3:**
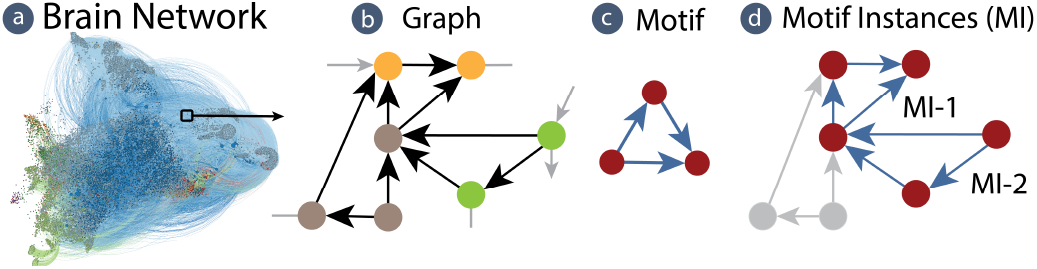
Motifs in brain networks. (a) Large connectomes like the FlyEM hemibrain dataset [44] can be interpreted as (b) directed graphs, where neurons represent nodes and synapses are represented as edges. (c) Network motifs are recurrent subgraphs in the larger brain network. For example, the graph shown in (d) contains two instances (MI-1, MI-2) of a feedforward motif (c).

**Fig. 4:**
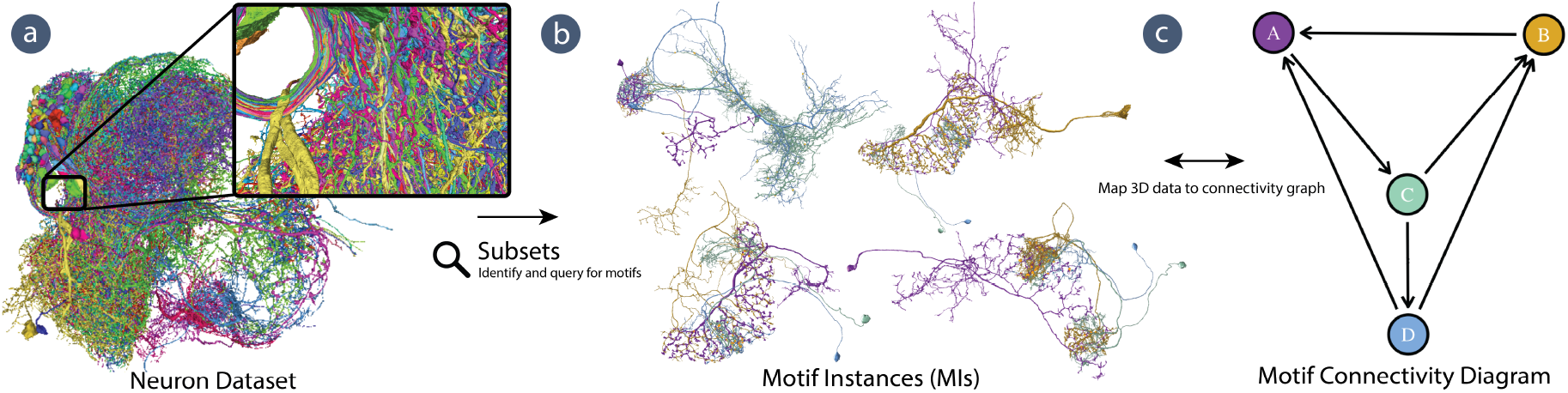
Data Overview. (a) The FlyEM hemibrain [44] dataset contains more than 21,000 neurons, forming a vast connectivity network. Here, we show 1% of those neurons as a Neuroglancer [30] rendering. An interactive version is accessible online [29]. (b) Small subsets of the dataset form specific network motifs. Each set of neurons forming a specific motif is called a motif instance (MI). (c) However, understanding motif connectivity in the complex 3D morphology of neurons is not trivial. Consequently, domain experts require more advanced tools to better understand how connectivity motifs are manifested in a set of neurons.

### Brain networks and motifs

Connectomics datasets with tens or hundreds of thousands of neurons can be interpreted as large *directed graphs*, with neurons as nodes and synapses forming edges between the nodes. Recent advances in microscopy and computational neuroscience have revealed characteristic non-random patterns in neural networks of invertebrate and mammalian brains, especially in the cerebral cortex. These small recurring wiring patterns [35], termed *network motifs*, suggest the hypothesis that neuronal connections are arranged using some basic building blocks with hierarchical order [52] (see Fig. 3).

### Connectomics Data

Analyzing neuronal tissue at the level of synapses requires ultra-high-resolution imaging techniques such as serial-section scanning electron microscopy (SEM), which can produce more than 1 petabyte of data for each imaged cubic millimeter [50] of tissue. In this paper, we demonstrate our visual motif query and analysis approach on the FlyEM hemibrain, one of the largest open-source connectomic datasets. It contains the entire central region of the fruit fly brain, including 21, 000 proofread neurons classified into more than 5, 000 types, over 20 million synapses, 13 main brain regions, and over 200 sub-brain regions [44]. This dataset is a major resource in the Drosophila neuroscience community and has led to exciting scientific discoveries [24, 27, 46]. *Vimo* uses the connectivity graph, 3D neuronal skeletons, and synapse locations of the hemibrain data (see Sec. 10).

## 4 Goal & Task Analysis

The idea of *Vimo* originated from meetings with neuroscience collaborators, who were excited about new EM datasets, but needed the means to analyze synapse-level connectivity. They want to identify and explore the morphological properties of motifs and synaptic chains without being overwhelmed by the 3D structure of intertwined neurons. At the same time, they need to see motifs in the original 3D space to understand the spatial relation of neurons and synapses.

Following the design study approach by Sedlmair et al. [49], we identified a set of domain goals and tasks in semi-structured interviews with five experts from the Harvard Center for Brain Science, HHMI Janelia, and Columbia University. All scientists are internationally recognized experts in analyzing neuronal circuits reconstructed from EM images. Three are experts in Drosophila connectomics. We also held unstructured interviews at an international connectomics conference, where we presented an early prototype to refine our goals and tasks.

### 4.1 Domain Goals

The neuroscientists’ main objective is finding small biologically relevant motifs in large neuronal networks. Further, they want to analyze interesting motif instances visually in more detail to fully understand how the motif’s connectivity relates to each neuron’s morphology.

#### G1 - Motif identification in large brain connectivity data

Our collaborators want to search large brain networks for instances of a wide range of small motifs (*≤*8 neurons) based on **neuronal connectivity** and additional **biological constraints** on the involved neurons and synapses. Neuroscientists want to specify details such as the *brain region* a neuron trajects, the *neuron type*, or the *number of synapses* a neuron makes in a specific brain region. For example, a specific type of ring neuron in the fly brain is involved in action selection tasks [4], and scientists want to analyze motifs involving this specific neuron type. Biological constraints ensure the expressiveness of motif queries when studying such behaviors. Furthermore, neuroscientists are interested if a specific motif is exceptionally over or underrepresented in the connectome network.

#### G2 - Analysis of motif instances

Since a motif search in a large brain network might result in dozens to thousands of hits, our collaborators need to be able to easily **identify a small set of interesting motif instances (i.e**., ***MIs*)** and then analyze them in high-resolution. Neurons form complex 3D shapes, span long distances, and connect with up to thousands of other neurons. Therefore, our collaborators previously struggled to **identify how a set of neurons forms a motif** and how the abstract connectivity motif relates to the actual 3D anatomy of the MI (see Fig. 4). However, this understanding is crucial for following information flow along neurons. In particular, our collaborators have questions such as *“Which branches and subbranches of the neuron make up the motif?”, “Are several spatially distinct parts of a neuron involved in the same motif?”* (see Fig. 4bc), or *“What is the anatomical context of involved neurons (e*.*g*., *in which brain regions are they located)?”* (see Fig. 5c).

**Fig. 5:**
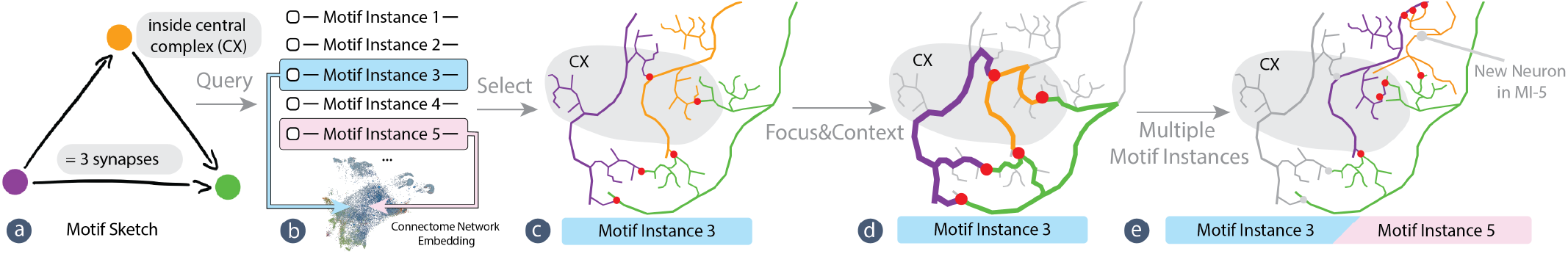
*Vimo* workflow. (a) Users query for motifs based on a sketch and set domain-specific biological constraints, like connectivity strengths and anatomical regions of interest, for targeted interrogation of the brain network. (b) After querying the connectome for a motif, users select found MIs for further investigation. (c) While exploring motif neurons and synapses (red dots), (d) users can adjust the visualization to gradually highlight the connectivity of neurons using our focus&context technique. (d) Next, users can continue to explore multiple connected MIs.

#### G3 - Identification and analysis of motif chains

In addition to exploring single MIs, our collaborators are interested in exploring motif chains. Starting from a motif of interest and a seed neuron, they want to **explore if the neuron is involved in different instances of the same motif**. This allows scientists to investigate if a single neuron computes similar functions, even in multiple MIs. Furthermore, our collaborators are interested in motif-based synaptic chains. Starting from a seed MI, they want to **follow chains of computation** to see if the same motif is formed consecutively over several connected neurons (see Fig. 10a). This analysis can give more holistic insights into information flow (e.g., from visual input neurons to the central brain of Drosophila).

#### Scalability Considerations

To analyze recently published large connectomes [44, 50, 62] with the aforementioned goals, *scalability* is the main requirement of our application. Those datasets contain tens of thousands of neurons and millions of synapses. Therefore, the entire analysis pipeline, including data access, algorithms, and interaction metaphors, need to support large data and scale to future datasets.

### 4.2 Tasks

Based on the above domain goals, we derived a set of analysis tasks *Vimo* needs to support:

#### T1 - Query for connectivity motifs

Scientists need a *fast* way to specify motifs and biological constraints for large graph queries (**G1**) without prior programming experience.

#### T2 - Identify biologically-relevant motif instances

Scientists need to be able to explore query results to identify interesting MIs (**G2**).

#### T3 - Explore the 3D structure of MIs

For an interesting MI, scientists want to explore the anatomy of its neurons and synapses in the context of the surrounding brain region (**G2**).

#### T4 - Visually identify the relation between neuron morphology and motif connectivity

After their initial exploration of an MI, scientists need to identify the parts of the neurons that make up the motif and analyze the detailed connectivity of the motif instance (**G2**).

#### T5 - Computationally and visually identify connected motif instances

Starting from the neurons of a selected motif instance, scientists want to explore whether the same neurons are involved in other instances of the same underlying motif (**G3**).

#### T6 - Trace synaptic chains starting from seed MI

Starting from a seed MI, scientists want to identify and visually follow synaptic chains made up of repeating specific motifs (**G3**).

## 5 *Vimo* Design and Workflow

Our main goal for *Vimo* is to provide neuroscientists with the means to easily explore the connectivity of connectomes and allow them to understand the network layout of neurons with their complex anatomical structure. Thus, we designed a workflow that offers intuitive user interactions and visualizations that highlight the connectivity of the data while keeping the anatomical data undistorted (see Fig. 5).

### Vimo Workflow

In collaboration with domain experts, we designed an *interactive visual query interface* based on sketching to search for network motifs in the data. The interface is aimed at domain experts with no programming experience and allows users to specify motif connectivity and biological constraints (**G1**). The main difficulty of connectivity analysis in high-resolution connectomic data is the complexity of both neuron morphology and connectivity. Neurons are densely packed in brain tissue and often highly intertwined. Synapses between two neurons might be clustered or spread out along the length of the neurons. Therefore, *Vimo* offers a *3D view* to display neurons and synapses of a selected MI in detail. This allows scientists to understand neuron shape and how motif neurons are intertwined. To support neuroscientists in the understanding of both morphology and connectivity, we designed a method for *gradually highlighting neuronal connectivity*, similar to continuous visual abstractions. Users can progressively focus on the underlying connectivity of the data while still retaining access to the morphology of neurons (**G2**). *Vimo* allows users to identify and follow interesting synaptic chains created by the repeated appearance of a motif. To reduce the search space for synaptic chains, we let users start with a seed motif or neuron, supporting user-driven data exploration (**G3**). Finally, throughout the design of *Vimo*, we focused on scalability by using datasets hosted in the cloud and only downloading small subsets to the user’s machine during runtime (see Sec. 10).

## 6 Interactive Motif Queries

In contrast to previous efforts in motif analysis that rely on programming languages [33, 38], *Vimo* provides a visual sketching interface to create queries. This gives scientists an intuitive approach to search for motifs. Additionally, *Vimo* provides real-time feedback on the number of occurrences of a sketched motif in the brain network to guide the user during the analysis process (see Fig. 6a).

**Fig. 6:**
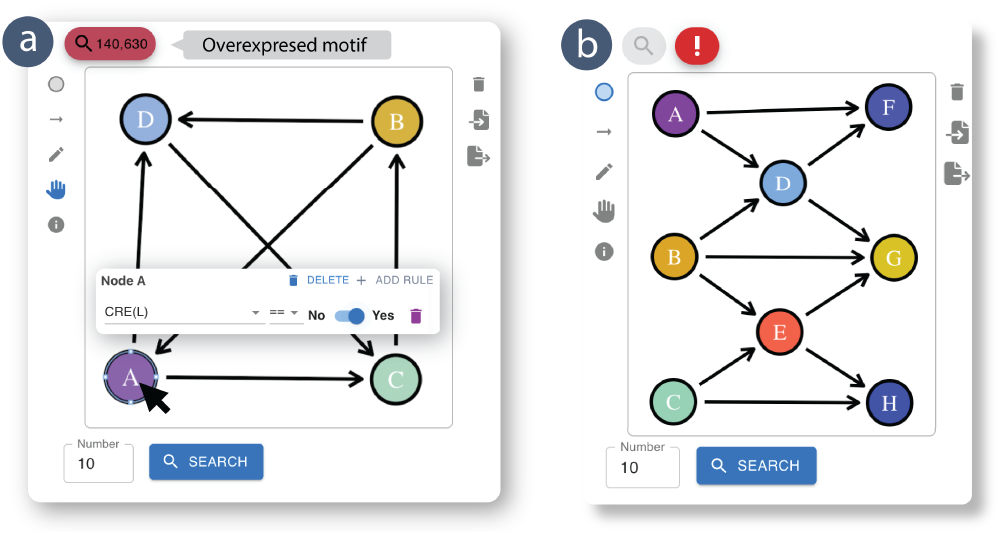
Motif sketching interface. (a) Users draw nodes and edges of a motif and define biological constraints on the nodes and edges. Additionally, *Vimo* gives real-time feedback on the absolute count of a sketched motif in the dataset and indicates whether a motif is over- or under-represented in the dataset (red label). (b) When searching the brain network for large motifs, *Vimo* warns users of potentially long runtimes using a red exclamation mark icon (b - top).

### 6.1 Motif Sketching

In *Vimo*, users search for motif instances by sketching an exemplar. They draw a set of nodes and edges to define the motif (i.e., neurons and the connections between them) (see Fig. 6), specify additional biological constraints, and inspect the results of their motif query.

### Defining Constraints

To create biologically meaningful queries, users can interactively set constraints on the nodes and edges of the motif sketch. Our query interface supports constraints on the *neuron’s type*, the *spatial location and brain region* of neurons and synapses, the *strength* of a connection, and the *neuron’s ID* (**T1**). For instance, when analyzing the hemibrain dataset, users can define one of over 5, 000 neuron types or trajectories through over 200 brain regions in the hemibrain dataset. We also introduce wildcard constraints that summarize certain families of neuron types to make queries more flexible. Scheffer et al. summarize all brain region constraints [43] and available neuron types [44]. *Vimo* offers an intuitive and expressive query builder interface to define one or multiple constraints per node and edge (see Fig. 6). Autocompletion of neuron types and brain regions further helps users select from thousands of available options.

### Guided Sketching

Deciding which motifs to study in greater detail is not always obvious to neuroscientists. During formative interviews, experts expressed interest in analyzing particularly common or rare motifs in the network. Therefore, as the user is sketching a new motif, we provide real-time feedback about the significance of the sketched motif. In particular, *Vimo* shows the number of occurrences of the motif in the brain network and whether the motif is estimated to be over- or under-expressed (**T2**). Counting the number of occurrences of a motif in a large graph in real-time is computationally infeasible. Therefore, we use precomputed counts of motifs with up to five nodes using a subgraph enumeration technique [32] that leverages a parallel version of the Kavosh algorithm [26] on a high-performance compute cluster. At run-time, we perform a simple look-up to display the number of occurrences of a sketched motif (see Fig. 6). Enumerating all occurrences of motifs with more than five nodes is computationally infeasible, as the number of possible motif configurations grows exponentially with the node count [32]. Nonetheless, *Vimo* supports finding representative MIs for motifs up to the size of 8 by limiting the number of MIs that are returned to the user (see Fig. 7). We also indicate the over- or underexpression of a motif in the brain network using a heuristic approach. We estimate a motif as over-expressed if its occurrence in the brain network is higher than in a random network. It is under-expressed if it occurs less frequently in the brain than in the random network. To analytically approximate the motif count in a random network, we build on a method [25] using Erdös models, a simple class of random graphs where each vertex-pair has a fixed probability of being connected by an edge. We indicate over- or under-expression with a red or blue badge in the sketching panel, respectively (see Fig. 6a).

**Fig. 7:**
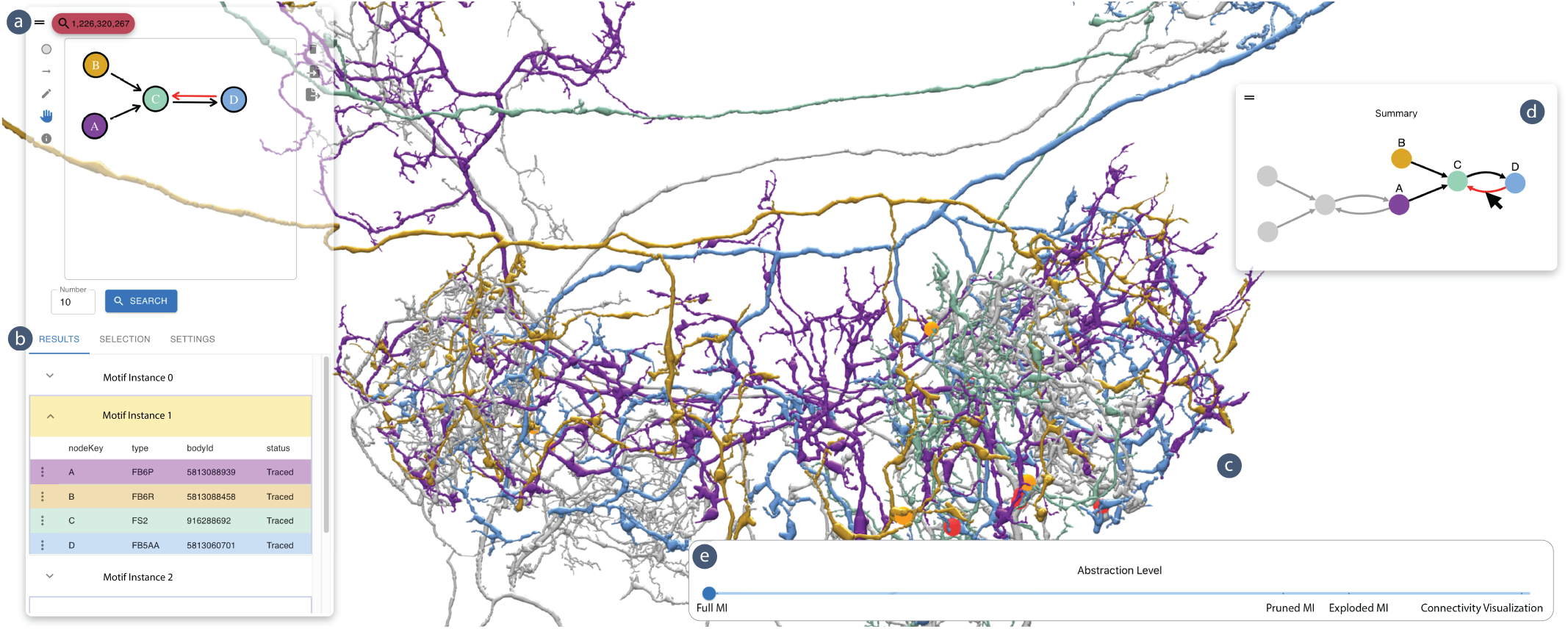
Overview of the *Vimo* user interface. (a) A motif sketch of a participant in the user study (see `motif-p2-pilot.json` at [57]) and (b) list view of resulting MIs. (c) The central view shows a non-simplified, full morphology 3D rendering of two MIs. (d) The multi-motif summary view visualizes the relationship between the selected MIs when following motif chains. (e) Users control the focus&context abstraction level using the interactive slider. Continuously simplifying the MI ensures preserving context between successive simplification steps.

### Running Motif Queries

*Vimo* automatically compiles the visual sketch into a *DotMotif* [33] string in real-time. Specifically, we convert a JSON representation of the motif sketch into a *DotMotif* query. Once users are satisfied with their sketch, we send the query string to a graph database to get a list of motif instances. Note that any graph database supporting *DotMotif* queries can be used. The graph database then returns a list of MIs fulfilling the properties of the sketch. This list is sorted based on the order in which the subgraph isomorphism search algorithm detects MIs. See Sec. 6.2 for the scalability of motif queries.

### Inspecting Motif Query Results

*Vimo* displays the results as a set of MIs in a list view (see Fig. 7b). To guide users in selecting a motif instance for further analysis, we show summarizing features for each neuron in an MI. In particular, we display the neuron’s type, ID, and proofreading status (**T2**). Clicking on an MI shows 3D models of the neurons and their synapses in the main view of *Vimo* (see Fig. 7c).

### Reproducing and Sharing Sketches

Users can import and export motif sketches as JSON files. This helps researchers reproduce their motif queries at later times and facilitates sharing of interesting discoveries with colleagues. A set of interesting motif sketches from the case study is available online [57] and in the supplementary material.

### 6.2 Large Motif Queries

*Vimo*’s motif sketching and interactive motif simplification scale to large motif instances (see Fig. 6b). However, subgraph isomorphism searches in large graphs are computationally expensive [32, 39], which becomes a factor as the number of motif nodes increases. For example, verifying the existence of a motif in a larger network is an NP-complete problem [10]. We use two strategies to limit the computational complexity of queries to maintain runtimes suitable for interactive systems. First, users can set targeted *biological constraints* on the nodes and edges, reducing the search space significantly. For example, neuroscientists are interested in motifs involving external ring neuron *(ExR1)* types. The hemibrain dataset [44] contains four neurons of type *ExR1*, which drastically reduces the search space by only querying the neighbors of those four *ExR1* neurons. Second, users can *limit the number of MIs* returned by a query (see Fig. 7b). Keeping this number low allows the search algorithm to terminate early without finding all MIs in the network. Visually inspecting a small number of MIs is often satisfactory for exploratory or initial analysis. We use a heuristic on the number of nodes in the motif (e.g., *≥* 5 nodes) to estimate if a motif query will run longer than 20 seconds, in which case we warn the user with an exclamation mark icon in the sketching interface (see Fig. 6b).

## 7 Motif Visualizations

Most connectome analyses require a detailed understanding of neuron wiring. However, due to the complex morphology of neurons, it is difficult to relate the motif connectivity to a 3D visualization of the neurons (see Fig. 4bc). Neither node-link diagrams nor planar visual encodings can accurately show spatial relationships between neurons (**T3**). 2D projections of neurons abstract their spatial entanglement, which is relevant to understand connectivity. Thus, we designed a focus&context method that gradually emphasizes motif connectivity in the 3D representation by pruning unimportant neuron branches and drawing visual links between connected neurons (**T4**) (see Fig. 8).

**Fig. 8:**
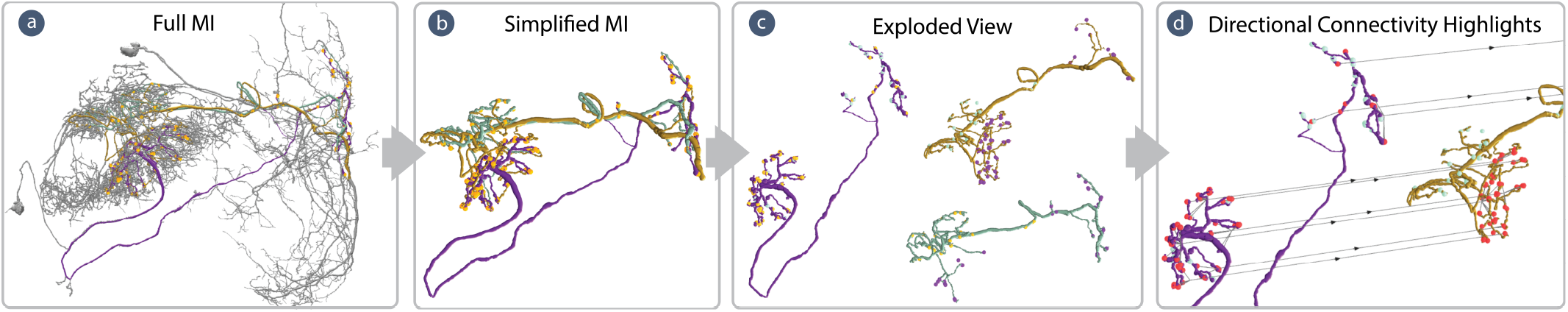
Gradual Motif Highlighting. (a) In *Vimo*, users preview a simplified version of the MI and maintain spatial context, as branches set for pruning are colored in grey. (b) Once neurons have been fully pruned, (c) users explode the view using a slider to decrease the visual complexity caused by tangled neurons. (d) In the exploded view, *Vimo* progressively groups synapses and creates bundled links between pre- and post-synaptic sites, minimizing visual clutter. Distinct link bundles signify separate clusters of synapses between the purple and yellow neurons.

### 7.1 Spatial Exploration

First, *Vimo* supports interactive inspection of the 3D spatial morphology of all neurons of a motif instance (**T3**). We render neurons based on their skeletons to ensure scalability. Skeletons are simplified stick-figure representations of neurons and are faster to download during runtime than meshes, as they require less data while retaining essential morphological details. For each skeleton node, *Vimo* uses the neuron’s diameter to adjust the thickness of the rendered tube, leading to more accurate visual representations. To allow users to quickly identify a neuron’s role in the motif, we color each neuron based on its color in the motif sketch (see Fig. 7). For context and to enhance a user’s spatial orientation, the 3D view in *Vimo* can show the outlines of 200 Drosophila brain regions, which are stored in Neuprint datasets [38].

### 7.2 Gradual Motif Highlighting and Abstraction

Our visual abstraction approach contains three main steps, all aimed at gradually highlighting motif connectivity, to help users understand how neurons form a motif (**T4**). Users can gradually move between the steps by dragging a slider (see Fig. 7e), which results in a continuous animation from one highlighting-and-abstraction level to the next.

#### Neuron pruning

First, we highlight motif connectivity by visually removing parts of a neuron that are not involved in the connectivity motif. Users can gradually peel away all parts of the neuron that are not in the motif, which reduces visual clutter and helps the user focus on the important parts of a neuron in relation to the motif (see Fig. 9). For each neuron in a motif instance, we start with a 3D skeleton representation (see Fig. 8a and Fig. 9a), which is interpreted as an undirected graph *N* = (*V, E*) with vertices *V* and edges *E*. First, we identify the set of skeleton vertices *P⊂ V* which will remain even after the neuron is fully pruned (see Fig. 9c). The set *S⊂ V* represents the spatial locations of motif synapses and is computed by mapping each synapse location to its closest vertex in *V*. Synapse locations are assumed to be part of the dataset. *P* is formally defined as

**Fig. 9:**
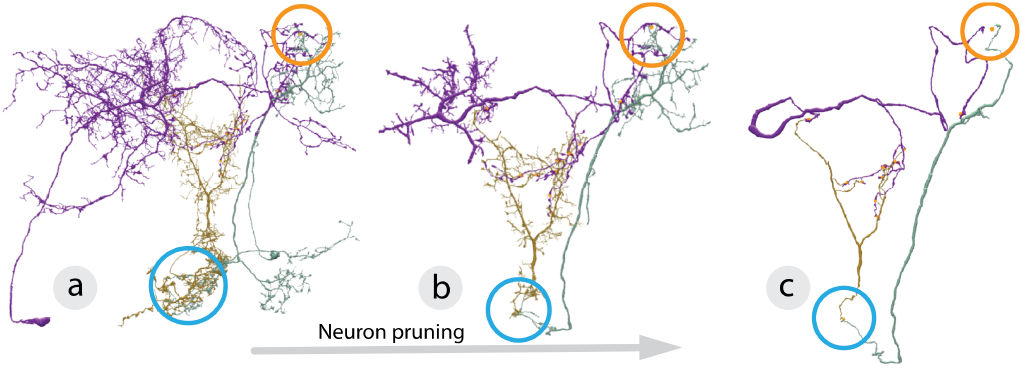
Motif pruning. *Vimo* gradually emphasizes neuron connectivity in MIs. We first show the full 3D morphology (a) and subsequently prune all branches unrelated to the motif (b) until only essential branches remain (c). In the process, neuron connectivity becomes visually more prominent (colored circles).

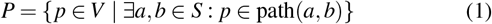

In other words, a vertex *p∈ V* is in *P* if there exists a path between any two motif synapses *a* and *b* that contains *p. P* represents neuron branches with synapses involved in the motif (**T4**). Next, we compute a distance value *d* for each vertex *v∈ V* to *P*, by calculating the geodesic distance *g* to its closest neighbor in *P*:

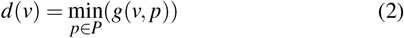

This allows us to gradually prune non-motif branches based on *d* by moving a slider until only branches with motif synapses remain (see Fig. 9bc). Vertices with higher distance values *d* are pruned earlier than vertices with smaller values. After an MI is fully pruned, the user sees the essence of all neurons making a motif (**T4**) (see Fig. 9c).

#### Exploded view

In the second step, we further reduce the visual complexity resulting from entangled neurons. We spatially pull all neurons apart into an exploded view after all non-motif branches are pruned (see Fig. 8c). This allows scientists to study the morphology and branching patterns of individual neurons and better visualize where synapses involved in a motif are located. To evenly distribute the neurons in space, we use the Saff and Kuijlaars algorithm [42] to compute the directions of the explosion. The algorithm evenly distributes *n* points on a unit sphere. We then compute an explosion direction for each neuron by calculating a vector from the sphere’s center to one of the *n* sampled points. In the exploded view, all pre- and post-synaptic sites between motif neurons are marked with spheres colored like their synaptic partner neuron (see Fig. 8c). This helps the user quickly grasp the neurons’ connectivity, even as the neurons are spatially apart (**T4**).

#### Connectivity Visualization

In the third step, we highlight connectivity in the exploded view inspired by context-preserving visual links [53]. We initially draw lines between the pre- and post-synaptic sites of the neurons (see Fig. 1e and Fig. 8d). We use two strategies to highlight important connectivity features and avoid visual clutter. First, we use hierarchical clustering of synapses and 3D bundling to aggregate lines. We chose hierarchical bundling as it can visualize different levels of synapse clusters and how they are distributed along the neurons. Based on feedback from our collaborators, we cluster synapses based on their spatial proximity to the neuron and which neuron they connect to. The user can set the bundling strength via the focus&context slider. As an alternative to the clustering approach, the user can also decide to only view connecting lines for user-selected synapses.

#### Design Alternatives

In an initial version of *Vimo*, we gradually moved from the pruned neuron view all the way to a node-link view by abstracting neurons into nodes and collapsing synapses to links. We hoped this would help scientists better visualize the relationship between the motif and the involved neurons. However, we ultimately abandoned this idea since scientists could already see the abstract motif in the sketching interface and had no use for such a simplified view. Additionally, based on feedback from visualization experts, we implemented a depth of field (DOF) effect to improve depth perception. However, during user testing, we found that DOF decreased users’ ability to follow long traces of neurons, as branches outside the focal plane are blurred. Therefore, users frequently adjusted the focused region to analyze an entire motif. Thus, we did not use the DOF effect during the evaluation.

## 8 Motif-based Synaptic Chains

Examining an individual motif instance (MI) offers a limited perspective on the connectivity of the associated neurons. To investigate how an MI is integrated within a broader network, researchers must visualize multiple interconnected MIs (**T5, T6**) (see Fig. 10). For example, when exploring pathways from visual input neurons to the central complex in Drosophila, conducting a multi-motif analysis proves to be pertinent.

### 8.1 Multi-Motif Analysis

*Vimo* supports the analysis of synaptic chains in two ways:

#### Neuron-centric analysis

First, domain experts are interested if a neuron forms the same motif with multiple partners (**T5**). For instance, in Fig. 10a, neuron A participates in two motif instances (MI-1, MI-2). Users can specify a *seed neuron* in the motif sketching interface by setting a neuron ID as a constraint to ensure that only motif instances involving that particular neuron are returned.

#### Synaptic pathway analysis

Second, neuroscientists are interested in following synaptic pathways that repeatedly include a sketched motif. For example, MI-1, MI-2, and MI-3 in Fig. 10 form such a synaptic pathway. To query for those, users start from a seed motif and set the ID of a sink neuron in the motif instance to a source neuron in the motif sketch through a context menu (**T6**). This approach facilitates the continuous construction of motif-based synaptic pathways downstream or upstream from a seed motif instance. Additionally, multi-motif analysis serves as a proxy for examining large motifs comprising numerous neurons. When it is computationally infeasible to search a brain network for instances of a large motif, users can alternatively construct the large motif iteratively by using smaller motifs as building blocks.

### 8.2 Multi-Motif Visualization

*Vimo* uses three strategies to visualize multiple MIs.

#### Connectivity Summary

*Vimo* provides an abstract overview of the connectivity of all selected MIs in a small node-link diagram (see Fig. 7d). The nodes and edges of the focused MI are colored, while all other elements are grayed out. This overview can help the user to stay oriented in the 3D view (**T5**). Selecting an edge in the summary view also highlights all corresponding synapses in red in the 3D view and the corresponding edge in the sketch panel (see Fig. 10).

#### Highlighting motif instances

Inspecting even one MI in 3D can be overwhelming due to the complex morphology of neurons. *Vimo* always focuses on a single MI, while all other unrelated neurons are grayed out to highlight the neurons of the MI in focus (**T6**). Users can quickly switch focus by clicking on a different MI in the list view.

#### Multi motif abstraction

We designed the continuous focus&context approach (see Sec. 7.2) to scale to multiple MIs. For each MI that a neuron participates in {*MI*_1_, …, *MI*_*n*_}, our pruning technique computes a set of distance values *D*(*v*) = {*d*_1_(*v*), …, *d*_*n*_(*v*) }for each vertex *v ∈ V* using Equation 2 (see Sec. 7.2). Thus, each vertex is assigned the minimum computed distance to avoid over-pruning an MI.

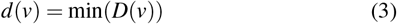

Pruning and exploding multiple MIs simultaneously helps experts to better understand the wiring of a synaptic chain (**T5, T6**) (see Fig. 10e).

## 9 User Interactions

We offer several interactions for visual motif analysis.

### Focus&context slider

An interactive slider controls the continuous focus&context approach for motif connectivity highlighting (see Fig. 7e) to maintain spatial context between consecutive views. First, dragging the slider starts pruning the MI continuously. After the MI is fully pruned, moving the slider further explodes the neurons in the 3D view. Finally, in the exploded view, users can control the bundling strength of the connectivity links using the slider (see Fig. 1e). For more details about our focus&context approach, see Sec. 7.2.

### Synapse highlighting

In *Vimo*, synapses are drawn as small spheres between the pre- and post-synaptic sites of the connected neurons. Visually highlighting all synapses between two neurons provides a simple yet effective visual clue to study where neurons connect (**T4**). A set of synapses connecting a pair of neurons is highlighted by either clicking on a single synapse sphere in the 3D view or by selecting an edge in the sketching panel or summary view (see Fig. 7d).

### Showing brain regions

Interactively rendering the outlines of anatomical brain regions (e.g., Central Complex in Drosophila) provides orientation and allows users to verify if a selected MI adheres to the constraints specified in the sketch **(T3)**. We show semi-transparent surfaces of the brain regions overlayed to the 3D renderings of the neurons. Users can interactively select which brain regions to show and quickly enable and disable the renderings.

### Graying out non-motif branches

While neuron pruning (see Sec. 7.2) helps remove unrelated branches to an MI, it removes context about the motif neurons that might be necessary during analysis. Therefore, *Vimo* allows graying out all non-motif branches of an MI (see Fig. 8a) to highlight important parts of the neurons and show the context of all other neuron branches by toggling a checkbox in the UI (**T3, T4**).

### Further Analysis

Once scientists identify interesting motif characteristics in their exploratory analysis, they need to perform an in-depth analysis on a set of MIs. This involves statistical analyses, such as studying the distribution of all synapses on a neuron (not just motif synapses). *Vimo* supports this step by integrating tightly with neuPrint [38], which provides a set of tools for general neuron analysis. Users can access neuPrint data for all neurons of an MI by using the context menu.

## 10 Data and Implementation

### Data

*Vimo* requires a connectivity network and proofread reconstructions of neurons in the form of 3D skeletons and synapses, including their spatial locations and pre- and post-synaptic partners. Meta-data, like neuron types and anatomical brain regions, are used to increase the expressivity of motif queries but are not required. Precomputed motif counts can guide motif sketching (see Sec. 6.1). All other computations can be performed during runtime. *Vimo* expects data in the neuPrint format [38]. neuPrint currently hosts five datasets (three hemibrain versions [44], the fib19:v1.0, and the MANC dataset [54]). New datasets are expected on the same platform soon. For the development and evaluation of this project, we used the *hemibrain v1*.*2*.*1* dataset, which includes 21, 000 proofread neurons and 20 million synapses.

### Scalability

Interactively exploring tera- and petascale connectome datasets requires scalable tool architectures. We leverage different data representations of neuronal circuits with varying memory requirements to achieve scalability. For instance, the electron microscopy imaging data of the hemibrain dataset is 26 TB in size [44]. In contrast, the skeleton representations of the reconstructed neurons require only *∼* 10 GB, and the pure connectivity graph compresses the data to *∼* 25 MB [44]. As a result, the data size required for certain analyses is reduced by a factor of a million by transforming the imaging data into a connectivity network. We exploit the different memory requirements by carefully choosing the data representation for each step of the analysis workflow. For instance, *Vimo* queries the compact connectivity network for motifs. Based on the detected and selected MIs, *Vimo* downloads a small set of the neuronal skeletons and synapses from the neuPrint server [38]. *Vimo* then uses an optimized web-based renderer to display those neuronal skeletons interactively. We only send skeleton vertices and edges to the GPU and use image-based rendering to generate visual primitives, like spheres and cones, in the fragment shader. Thus, we avoid costly data transfers between the CPU and the GPU. Hence, *Vimo* can be used on consumer-level hardware without RAM and GPU requirements.

### Implementation

*Vimo* is implemented as a web application using a React-based client and a Python-based server. *Vimo* builds on software libraries like *NaVis* [45] for data processing and a modified version of the WebGL-based *SharkViewer* [61] for 3D rendering. The hemibrain dataset [44] is hosted remotely in the *neuPrint* ecosytem [38]. Motif sketching is implemented using the *Paper*.*js* library. *Vimo* translates the visual sketch to an optimized *Cypher* query [17] using the *Dotmotif* [33] language as an intermediate step. This query is sent to a remote graph database to determine a list of MIs. *Vimo* is open-source [57], and we offer instructions for users in our tutorials. We also have released a domain-agnostic React component [9] for motif sketching to lower the entry threshold for motif analysis across disciplines.

## 11 Evaluation

We report on an in-depth case study and qualitative user study to evaluate the usability and usefulness of *Vimo*.

### Participants

We evaluated *Vimo* with 7 experts (P1 - P7, 3 male, 4 female) from the Harvard Center for Brain Sciences and HHMI Janelia. Two participants are also co-authors. To limit participants’ time commitment, three participants performed the case study, and the remaining four participants completed the user study. All participants are experts in analyzing neuronal circuits reconstructed from EM image data, and four are experts in Drosophila connectomics (6 postdoctoral researchers and 1 Ph.D. student in neuroscience). None of the participants currently use interactive tools for motif analysis, even though 6 out of 7 participants rated motif analysis as important for their research.

### Setup

We met with each participant for 90 minutes in person or on a Zoom video call. After a short introduction to the tool, all users, who had no hands-on experience with *Vimo*, steered the tool themselves. We asked all participants to think out loud to capture their thoughts.

#### 11.1 Case Study: Exploratory Analysis

We report on an exploratory case study with three domain experts specializing in analyzing neuronal circuits in the Drosophila brain. We provided no specific tasks, as we wanted to test what types of analyses an experienced neuroscientist would conduct. We describe two motifs in detail that were analyzed by the experts during the session.

##### Feedforward outputs of ExR neurons

First, the expert searched for a familiar connectivity motif to verify the tool’s reliability. They started by sketching a feed-forward motif involving an external ring (*ExR*) neuron, an ellipsoidal body (*EB*) neuron, and a neuron without any constraints **(T1)**. The motif sketch can be reproduced by loading case-study-motif-1.json, available on our GitHub repository [57]. The expert iteratively refined their sketch by adding more constraints until they found an MI that matched the desired characteristics. Next, they selected an MI that included one *ExR* neuron and two *EPG* neurons (Ellipsoid body - Protocerebral bridge - Gall) **(T2)**. After inspecting the 3D renderings, they used the summary view to highlight synapses between the *ExR* neuron and both *EPG* neurons (see Fig. 7d). The expert indicated this helped confirm that the *ExR* neuron forms synapses with each *EPG* neuron on different arbors **(T3)**. They decided to study this observation in more detail using the focus&context slider and first pruned all unrelated motif branches of the neurons. Next, they iteratively moved back and forth between the non-exploded and exploded views to understand how the neurons entangle at areas of strong synaptic connectivity. Next, they studied the bundled lines between the pre-synaptic sites of the *ExR* neuron and one of the *EPG* neurons. They stated that neuron pruning and the exploded view helped them find a previously unknown connection and quickly identify which synapses might be considered biological noise **(T4)**. Finally, the expert was interested in performing a *neuron-centric multi-motif analysis* (Sec. 8.1) of the currently selected MI and other neurons expressing the same pattern. Specifically, they wanted to study if there are other MIs of the sketched feed-forward network that also involve the currently selected *ExR* neuron and one of the selected *EPG* neurons. Hence, they searched the query results for a second MI including these neurons, and added it to the 3D view **(T5)**. They frequently switched the focus between both MIs to better understand their spatial relationships and used the multi-motif abstraction (Sec. 8.2) to compare the connectivity of both MIs.

##### Circular connections in visual input neurons

Next, the expert sketched a circular motif of three visual input neurons, specifically tuberculo-bulbar (*TuBu*) neurons and ellipsoid body ring (*ER*) neurons **(T1)**. The motif sketch and the studied motif instance are available for reproducible results (`case-study-motif-2.json` and `case-study-motif-2-instance.json` at [57]). Based on the motif counts in the sketch panel, the expert found that circular connections are underrepresented in the dataset, making it interesting for detailed analysis **(T2)**. After analyzing an MI in 3D using *Vimo*’s focus & context technique, the expert identified that all synapses are clustered tightly at a specific location **(T3, T4)** (see Fig. 11ab). Based on this observation, the expert was interested if other neurons of the same type formed the same motif but expressed stronger connection strengths. Hence, they increased the synapse strength constraints in the sketch and found another MI with an even stronger synapse cluster close to the previously observed cluster **(T1, T2)**. As a final step, the expert used the exploded view to study the internal structure of the cluster and inspected the bundled lines to learn how the circular motif is expressed within the cluster **(T4)**. Overall, the expert stated that neuron pruning and the bundled connectivity links helped them quickly judge if synapses are clustered tightly or distributed randomly on motif neurons.

**Fig. 10:**
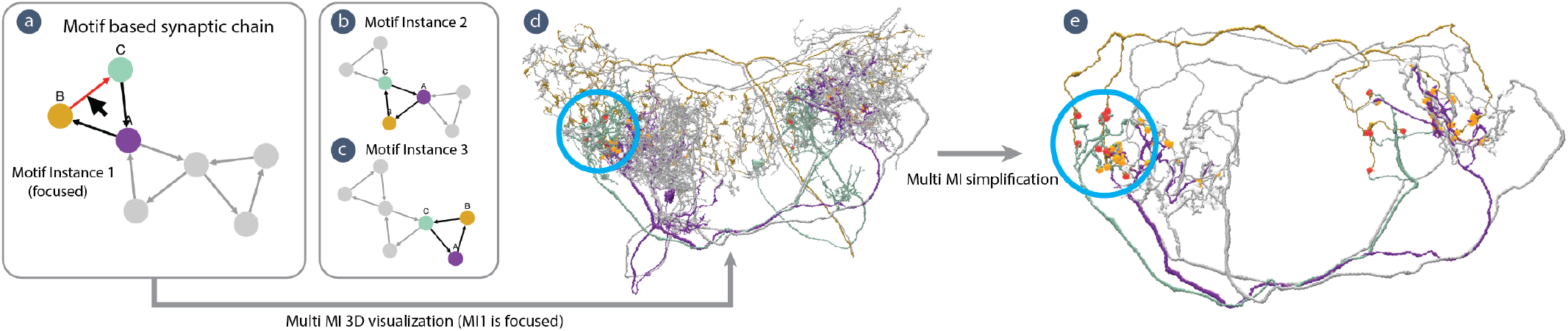
Visualizing multi-motifs. (a - c) *Vimo* supports exploring connections between multiple motif instances. For instance, the set of neurons in (a) form a circular connectivity motif three times. *Vimo* supports tracing such synaptic chains by repeatedly searching for instances of the same motif. (d) *Vimo* only highlights one MI at a time and defocuses all other MIs in grey. (e) Our focus&context technique scales to multiple MIs. By dragging the interactive slider (see Fig. 7e), we peel unrelated branches of all MIs to emphasize neuron connectivity (see blue circle).

**Fig. 11:**
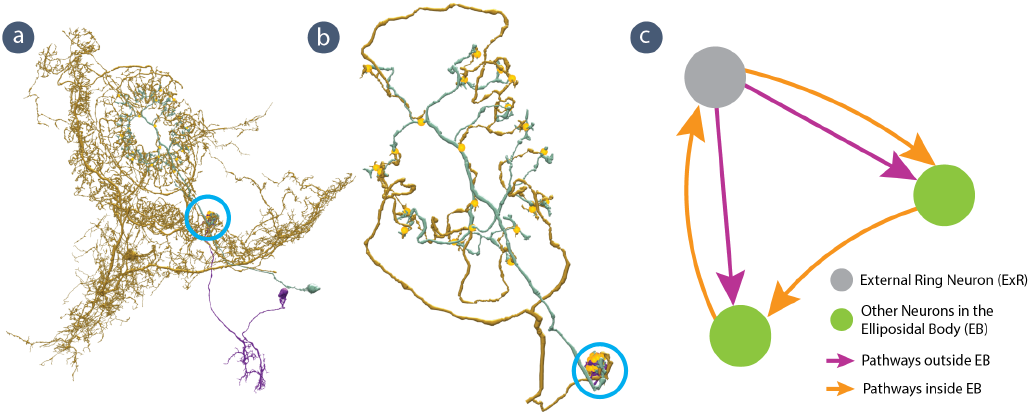
Vimo evaluation examples and tasks. (a) In the exploratory case study, a participant explored a circular connectivity motif between visual input neurons. By simplifying the motif using the focus&context technique, they could identify a synapse cluster; See the colored circle in (a) and (b). (c) In Task 1 of the qualitative user study, participants were asked to sketch this known biological motif between *ExR* neurons and other neurons in the ellipsoidal body (*EB*) of Drosophila [24].

### 11.2 Qualitative User Study

To evaluate the usability and usefulness of *Vimo* in a qualitative user study, we asked participants to perform two tasks: motif sketching and analyzing motif connectivity with our focus&context approach.

### 11.3 Task 1: Motif Sketching and Querying

We asked participants to sketch and query for a connectivity motif recently discovered (**T1**) [24]. The motif describes *ExR* neurons and other *EB* neurons that form connections inside and outside the *EB*. This motif contributes to the context-dependent action selection behavior of Drosophila. A correct motif sketch and related motif instance are available online to reproduce results [57]. We provided a schematic illustration of the motif and its constraints (see Fig. 11c) to all participants. Sketching the connectivity between the motif neurons was straightforward for all participants while defining node and edge constraints involved a learning curve. Two participants needed help from the study conductor to specify appropriate constraints. We observed that the performance of defining constraints depends on the participant’s familiarity with fly brain anatomy. Participants familiar with the dataset could easily identify how to translate the instruction *‘outside EB pathway’* into an edge constraint. Our survey results show that all four study participants found the motif sketching interface useful (see Fig. 12, Q2). However, for querying large motifs involving more neurons (e.g., *n≥* 6), only 3 out of 4 participants found motif sketching useful because of visual clutter in the sketching interface. During our pilot study, we found that neuron-type wildcards can improve the utility for defining node constraints (Sec. 6.1). Therefore, we added this functionality that was then widely used for the remainder of the user study. Additionally, participants suggested changing synapse strength constraints to a default of 10 synapses to avoid repetitive interactions.

**Fig. 12:**
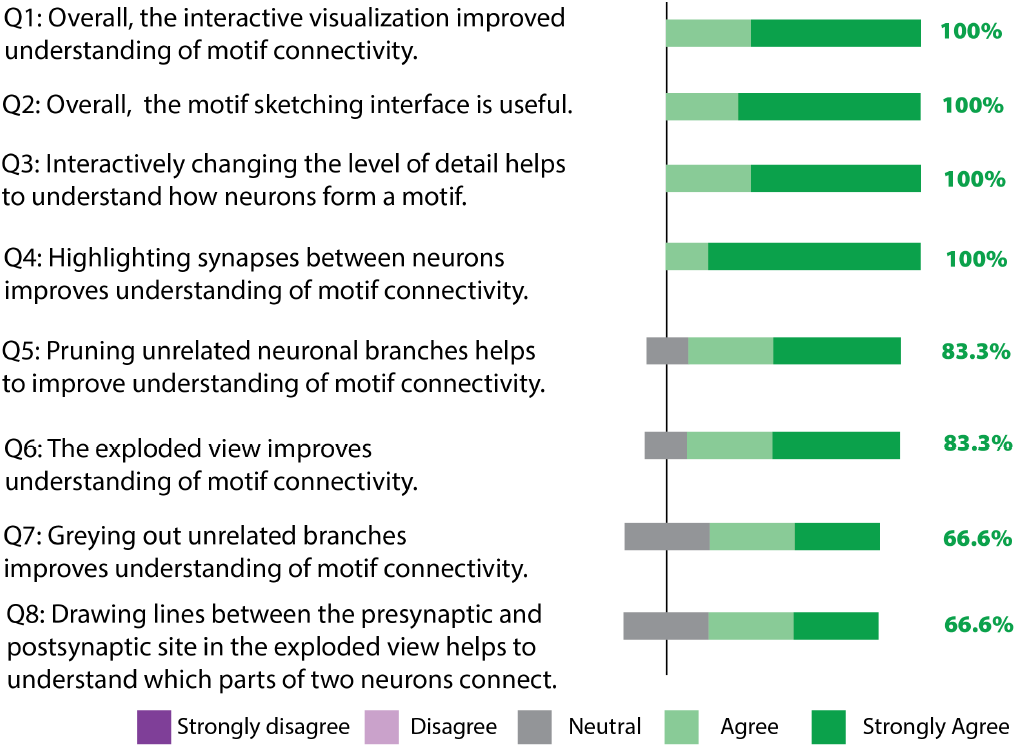
User ratings. We show neutral responses in gray, positive responses in green, and the percentage of users who (strongly) agreed.

### 11.4 Task 2: Analyzing Motif Connectivity in 3D Data

In the second task, we provided users with a motif and the 3D view of a motif instance and asked them to identify the main parts of the motif neurons connected in the 3D data (**T3, T4**). We tested two conditions: In the first condition, participants had access to the full functionality of *Vimo*, including our focus&context technique. In the second condition, participants could not use our focus&context method. We used two MIs describing feedback loops of CX neurons [57]. We counterbalanced between both conditions. Every participant had to perform the task with both conditions. To test their understanding of the motif connectivity in 3D, we asked participants to draw an illustration of the neuron branches involved in the motif and their connections for both conditions. We observed that participants frequently used the interactive slider (see Fig. 7e) to prune MIs maximally and to toggle between the non-exploded view and the exploded view to get a better view of neuron entanglement. Therefore, using our focus&context technique led to less cluttered illustrations by the participants, indicating a clearer and better understanding of motif connectivity due to interactive simplification. We show an example user illustration in the supplemental material. Additionally, we collected survey responses from all participants about our interactive motif abstraction approach (see Fig. 12), demonstrating that neuron pruning (**Q3, Q5**), synapse highlighting (**Q4**), and the exploded views (**Q6**) are among the most effective interactions of *Vimo*.

### 11.5 Key Findings

#### *Vimo* leads to fast, iterative analysis

We observed that users often queried a specific motif but forgot to specify certain constraints in the motif sketch. *Vimo* allows users to quickly iterate on their motif sketches and queries, leading to more relevant motif analyses.

#### Biological constraints are essential for motif analysis

We found biological constraints to be essential for expressive motif queries. Specifically, wildcards for neuron types were perceived well, constraining a neuron to a group of types. For instance, the *ExR\** type requires a neuron to be in any of the *ExR1* - *ExR8* types.

#### Exploded views improve spatial motif understanding

The gradual transition of neurons into the exploded view helped users understand how neurons entangle. Understanding the complex entanglement is crucial for analysis but also an obstacle for mentally mapping the motif graph structure to the 3D visualizations of neurons.

#### *Vimo* improves the state of the art

All study participants agreed that *Vimo* improves their motif analysis workflow. Users particularly liked the interactive motif sketching, neuron pruning, and synapse highlighting for their visual analysis in 3D (see Fig. 12, Q2, Q3, Q4).

## 12 Discussion

### User expertise for high-resolution connectivity analysis

It was challenging to find qualified experts for the user study. Many neuroscience labs are interested in connectivity analysis at the nanometer scale, however, few have actually done it. Datasets have only recently become available, and with a lack of scalable computational tools, many scientists have not yet started on this endeavor. We hope that *Vimo* can reduce the barrier of entry for this type of research and attract more labs and researchers to analyze high-resolution connectivity.

### Limitations

*Vimo* focuses on the interactive analysis of motifs and quick, iterative refinement of motif queries. Therefore, the strength of our tool is in analyzing relatively small motifs with a limited number of nodes (*n≤* 8). We determined this threshold through iterative testing. While our visualization approach scales to larger motifs, running large motif queries becomes computationally expensive. Such large queries potentially require hours of computation time, which would hinder the interactivity of our tool. Furthermore, while *Vimo* supports the visual analysis of multiple connected motif instances, large-scale comparisons of motif instances distributed across the entire dataset are not yet supported. Finally, in *Vimo*’s current design, researchers need a prior hypothesis about relevant motifs. In other types of analyses, researchers start from a fixed set of neurons and are interested in the set of expressed connectivity motifs. This exploratory approach is inverse to the workflow in *Vimo* and not yet supported.

### Tradeoff between accuracy and visual abstractions

A main design decision of our approach is not to distort neuron anatomy but rather enhance the user’s perception of the motif connectivity in the original data. Many previous approaches have focused on visual abstractions that simplify neuron anatomy [1, 36]. We take a complementary approach and focus on highlighting connectivity in the original data. This comes at the cost of a higher cognitive load than using simplified 2D representations. However, it allows scientists to better understand the detailed spatial make-up of an MI.

## 13 Conclusion and Future Work

*Vimo* takes a first step towards visual motif analysis for nanoscale brain data. The core idea of our approach is to enable quick and intuitive sketching of interesting motifs and to allow specifying biological constraints. We support a detailed motif analysis in the original 3D space, using a focus&context approach. We improve understanding of the data by gradually highlighting motif connectivity relative to the 3D structure and arrangement of neurons and synapses. *Vimo* is a key stepping stone towards neuronal pathway analysis at scale, where structural and functional data are combined for a better understanding of the brain. While *Vimo* is targeted to a specific audience, our motif sketching and novel simplification method for spatial graphs are domain agnostic. Specifically, the focus&context approach can, in principle, be adapted to any spatial graph data where nodes exhibit complex spatial morphology.

In the future, as more large-scale and proofread datasets become available [12–14,34,44,50,58], we want to extend *Vimo* to support data from other organisms, such as mice and humans. *Vimo* is currently being used at HHMI Janelia to analyze the next generation of connectome datasets. With these new datasets, inter-specimen comparative motif analysis will be within reach, posing new challenges for visualization. Thus, future tools should support analyses into whether similar neurons across specimens form similar motifs.

## 14 Acknowledgments

This work was supported by NSF awards IIS-1901030, NSF-IIS-2239688, NCS-FO-2124179, and R24MH114785. We gratefully acknowledge internal financial support from the Johns Hopkins University Applied Physics Laboratory’s Independent Research & Development (IR&D) Program for funding portions of this work. We also thank the participants of our user study and the anonymous reviewers for their valuable feedback.

