## Supplemental Figures for "Vimo: Visual Analysis of Neuronal Connectivity Motifs"

### ViMO - Visual Analysis of Neuronal Connectivity Motifs – Supplementary Material –

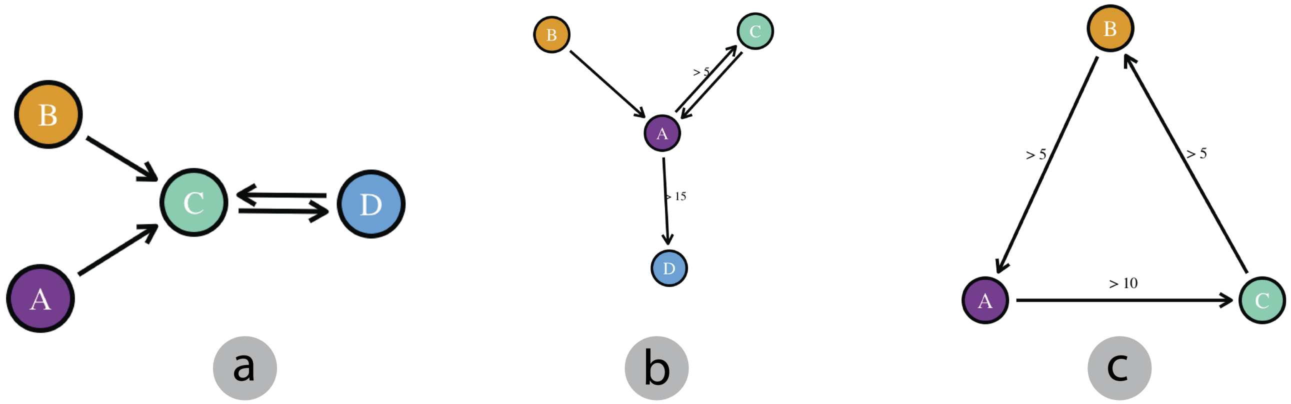


Fig. 1: **Examples of motifs sketched in the case study.** After sketching each motif, users successfully queried the brain network for MIs and analyzed MIs in 3D. To reproduce these motif sketches, users can import JSON files (a) `motif-p2-pilot.json`, (b) `motif-p1-study.json`, and (c) `case-study-motif-2.json` from https://github.com/jakobtroidl/neuronal-motifs/tree/main/example_motifs.

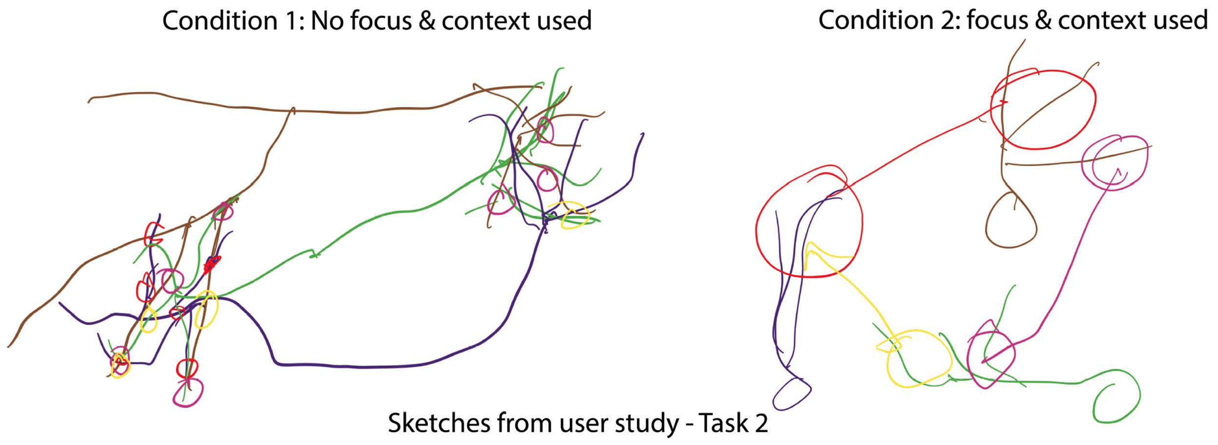


Fig. 2: **User illustrations of motif connectivity.** In Task 2 of the qualitative user study, experts illustrated their understanding of motif connectivity. For the first condition, they were not allowed to use Vimo's focus&context approach, while they were allowed to use it in the second condition. Using our focus&context method resulted in clearer illustrations, indicating a better understanding of motif connectivity.
